# *Candida glabrata YPK2* is a multidrug susceptibility locus

**DOI:** 10.64898/2026.05.15.725557

**Authors:** Lucia Simonicova, Thomas P. Conway, Thomas Krüger, Axel A. Brakhage, W. Scott Moye-Rowley

## Abstract

The biological conservation between fungi and mammals due to a common ancestor has made development of selective antifungal drugs a difficult challenge. Further complicating this situation is the selection of antifungal drug-resistant organisms during drug treatment. The pathogenic yeast *Nakaseomyces glabratus* (called here *Candida glabrata*) presents an especially challenging organism due to its tendency to frequently lose susceptibility to the major antifungal drug class the azoles. Additionally, *C. glabrata* develops resistance to echinocandin drugs, a second, more recently described antifungal agent at 10 times the rate of other organisms. Previous work has established that the sterol responsive transcriptional regulator Upc2A is a key determinant of azole susceptibility in *C. glabrata* and plays a role in echinocandin resistance. We used a biochemical approach to identify proteins that co-purified with Upc2A and identified the Ypk2 AGC kinase as an interacting protein. Strains lacking *YPK2* exhibited increased susceptibility to fluconazole and the echinocandin caspofungin. A *ypk2Δ* strain failed to normally induce transcription of several *ERG* genes but exhibited normal induction of the *CDR1* ATP-binding cassette transporter gene. Isogenic *ypk2Δ* strains were also highly susceptible to the three major classes of antifungal drugs, indicating that this kinase behaves as a multidrug susceptibility factor. RNA-seq analyses indicated that the transcriptional response to exposure is different for each drug and each response is differentially altered upon loss of Ypk2. Our data indicate that Ypk2 plays an important role in coordinating gene expression that impacts susceptibility to all major antifungal drug classes.

## Introduction

A threat to continued chemotherapeutic treatment of infectious microorganisms is the development of resistance to effective antibiotics. This problem is especially acute in pathogenic fungi for which only 3 commonly used types of antifungal drugs are available (FISHER *et al*. 2022). These drugs include the azoles, that block biosynthesis of the essential fungal sterol ergosterol, and the polyenes that bind through sequestering ergosterol in the plasma membrane and triggering membrane defects (ANDERSON *et al*. 2014). The most recently developed antifungal drugs are the echinocandins that are thought to act via inhibition of the β-glucan synthase enzyme, also located in the plasma membrane (Reviewed in (DENNING 2003)). Other antifungal drugs are under development but none are used to the extent of these three therapies.

*Nakaseomyces glabratus* (hereafter *Candida glabrata*) is a pathogenic yeast with a clinically problematic ability to develop robust antifungal drug resistance. This tendency is so high that current clinical guidelines recommend use of the echinocandin class of drugs rather than the orally available azole drugs (PAPPAS *et al*. 2016). *C. glabrata* is the second most common species associated with candidemia after *Candida albicans*, making this a pathogen of major significance (KIM *et al*. 2016; KATSIPOULAKI *et al*. 2024; CUGNATA *et al*. 2025).

Much of what is known explaining drug resistance in *C. glabrata* comes from the analysis of fluconazole-resistant isolates. A large number of clinical isolates have yielded the consistent finding of substitution mutations in a Zn_2_Cys_6_-DNA-binding domain-containing transcription factor called Pdr1 (VERMITSKY AND EDLIND 2004; TSAI *et al*. 2006; FERRARI *et al*. 2009). These gain-of-function (GOF) forms of Pdr1 behave as hyperactive transcriptional activators and drive high level expression of downstream target genes and strongly decreased azole susceptibility. One of the major Pdr1 target genes that contributes to azole resistance is the ATP-binding cassette transporter-encoding *CDR1* gene (TSAI *et al*. 2006; VERMITSKY *et al*. 2006). GOF variants of Pdr1 exhibit high constitutive levels of *CDR1* transcription with these increased levels required for the observed reduction in azole susceptibility. The Cdr1 ABC transporter is thought to act as an ATP-dependent drug efflux pump and lower levels of azole drugs in the cell (Reviewed in (PRASAD AND GOFFEAU 2012)).

While the Pdr1/*CDR1* circuit is important for azole susceptibility, the target of azole drugs is the lanosterol α-14 demethylase enzyme, encoded in *C. glabrata* by the *ERG11* gene (NAKAYAMA *et al*. 2001). Exposure of wild-type *C. glabrata* cells to azole drugs like fluconazole triggers enhanced transcription of *ERG11* through activation of a sterol-responsive transcription factor called Upc2A (NAGI *et al*. 2011). We discovered a transcriptional connection between Upc2A and both the *PDR1* and *CDR1* genes, consistent with the view that inhibition of ergosterol biosynthesis leads to induction of *ERG* gene expression but also transcription of the *PDR1* and *CDR1* genes (VU *et al*. 2019; VU *et al*. 2021). Disruption of the *UPC2A* gene prevents both activation of *ERG*genes in addition to *PDR1* and *CDR1*.

Control of the activity of Upc2A has been argued to be regulated at the level of nuclear localization of this factor by direct ergosterol binding (YANG *et al*. 2015). Low ergosterol levels are believed to cause a conformational change that releases Upc2A from a cytoplasmic tether and allow its import into the nucleus with attendant gene activation (TAN *et al*. 2022). To determine if other factors might act to modulate activity of Upc2A, we used a biochemical approach to identify proteins that co-purify with a tagged form of Upc2A. One of the proteins found in this approach was the AGC kinase Ypk2. This kinase, and its homologue Ypk1, has been extensively studied in *Saccharomyces cerevisiae* (See (ROELANTS *et al*. 2017) for a recent review). Most of the work on the *S. cerevisiae* Ypk1/2 kinases has focused on the important role these kinases play in control of biosynthesis of sphingolipids (BRESLOW *et al*. 2010; ROELANTS *et al*. 2011), a key class of membrane lipids. More recently, evidence has been provided that these kinases also control trafficking of the fungal sterol, ergosterol, in the plasma membrane (MURLEY *et al*. 2015; ROELANTS *et al*. 2018). Our data reveal that Ypk2 is required for normal expression control of genes involved in the biosynthetic pathway that produces ergosterol in *C. glabrata*. This effect is exerted at the level of activity of Upc2A. Strains lacking the *YPK2* gene exhibit increased susceptibility to fluconazole, amphotericin B and caspofungin, providing evidence that Ypk2 function is involved in the ability of a cell to respond to stresses exerted by the three major classes of antifungal drugs commonly used in clinical medicine.

## Materials and Methods

### Strains and growth conditions

All strains used in this study are listed in Table 1. *C. glabrata* strains were grown at 30°C overnight. Liquid overnight cultures were diluted to OD_600_ = 0.2 and continued to grow to the mid-log phase of OD_600_ =∼1. Rich YPD media (yeast extract 1%, peptone 2%, glucose 2%) were used for non-selective growth and drug treatments. Media were supplemented with 50 mg/mL of nourseothricin to select for strains carrying the *natMX4* gene conferring nourseothricin resistance on the plasmids or as a part of deletion/integration DNA cassettes. Synthetic complete media lacking histidine (0.67% yeast nitrogen base with ammonium sulphate, 2% glucose, complete supplement mix lacking histidine) were used for selective growth of *C. glabrata* strains transformed with *YPK2* deletion cassette containing the *HIS3MX6* selection marker.

**Table 1.**
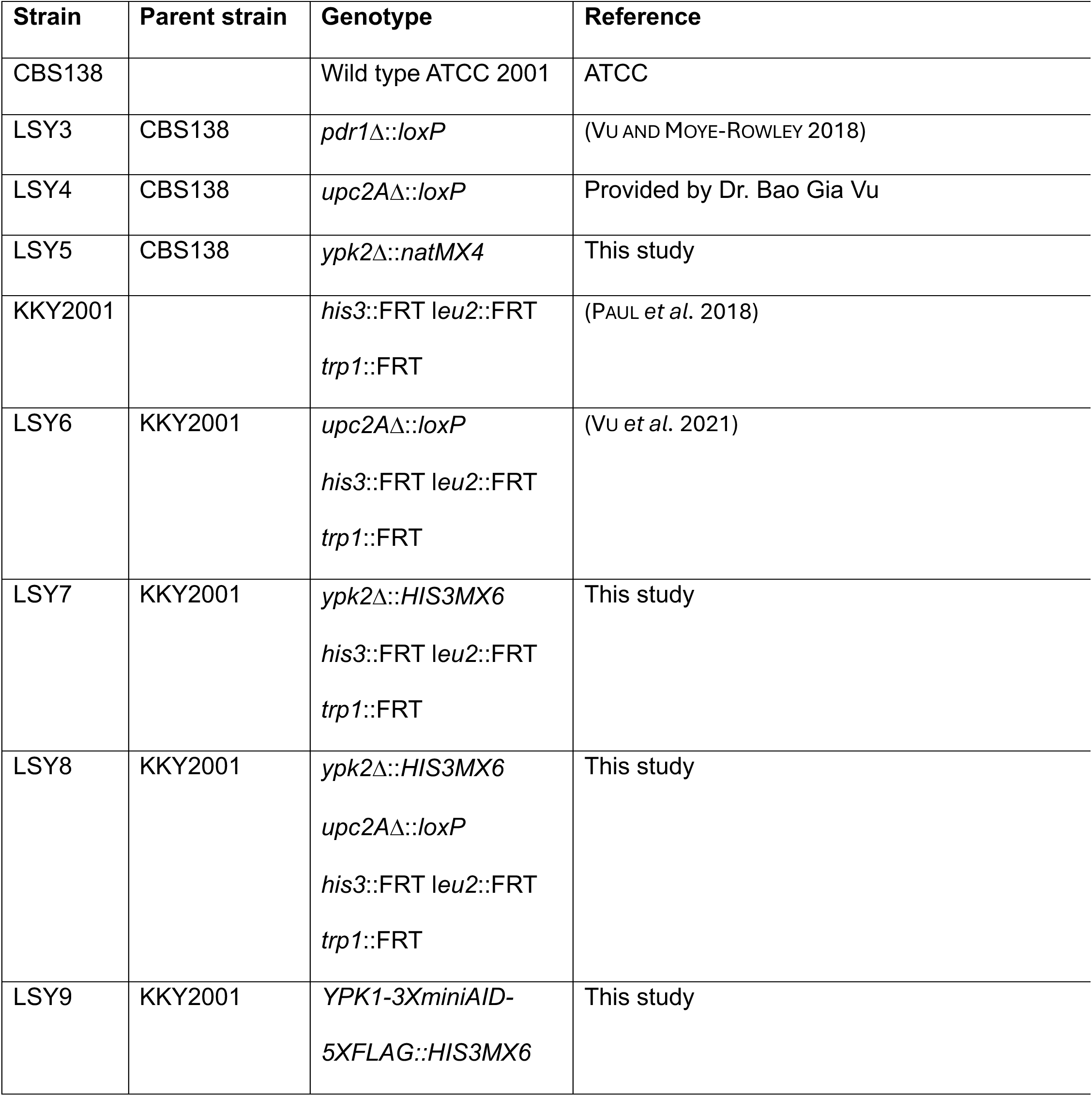
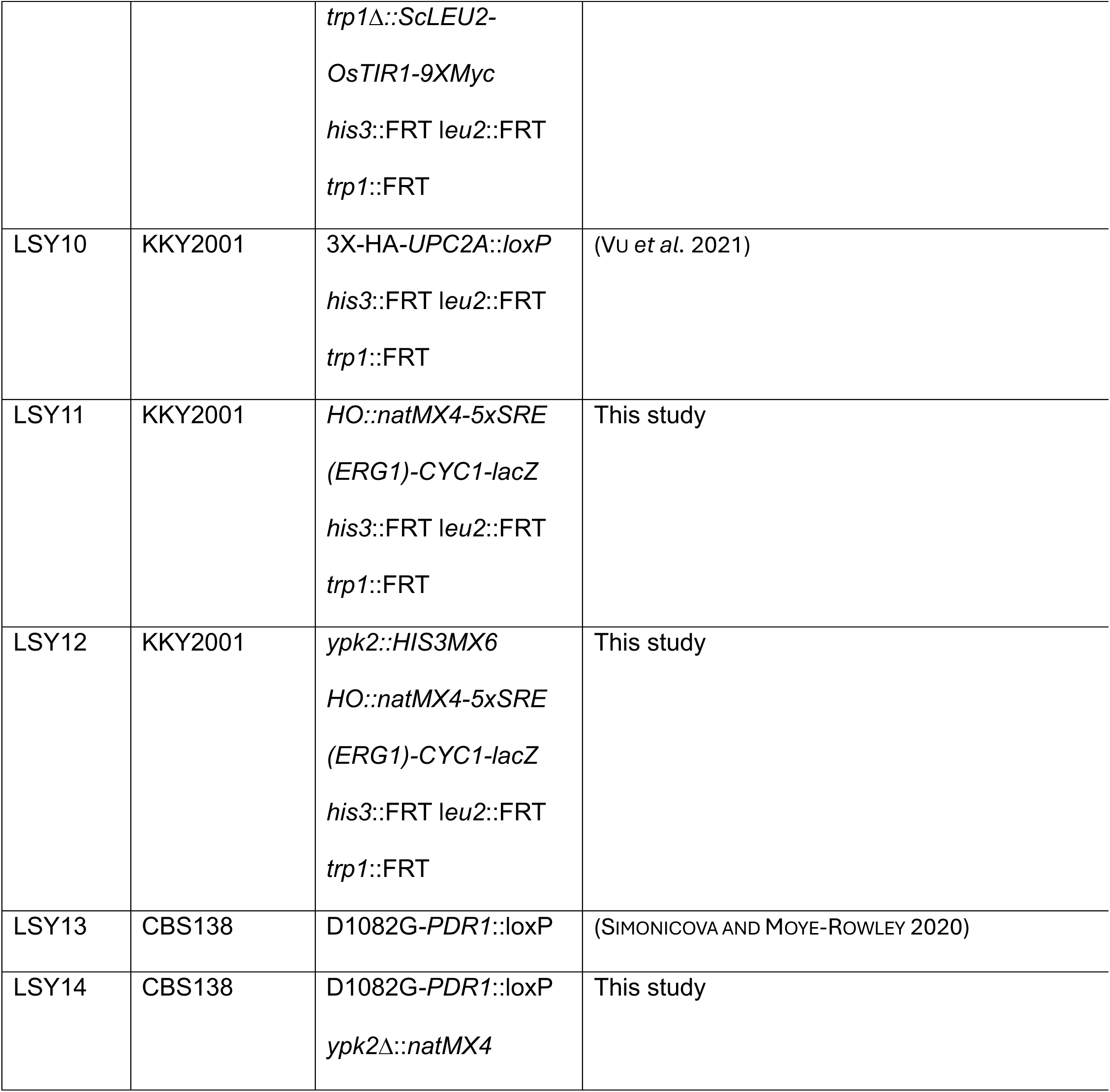
Strains used in this work.

### Plasmids

All plasmids used in this study are listed in Table 2. Cloning was done using Gibson Assembly Cloning kit (NEB) and the identity of recombinant vectors was confirmed by restriction enzyme analysis of plasmid DNA and by sequencing. To clone amino-terminally TAP-tagged *UPC2A* allele into the low copy *natMX4* vector, the *UPC2A* promoter and the *UPC2A* open reading frame (ORF) were PCR-amplified from the pLS20 and the N-terminal TAP tag was PCR-amplified from the pLS1. PCR fragments were cloned into the pLS20 that contained *UPC2A* terminator and was first linearized by SapI and SpeI to eliminate the *UPC2A* promoter and *UPC2A* ORF. N-terminal TAP-tag fragment was cloned upstream of the *UPC2A* ORF and downstream of the *UPC2A* promoter. To clone the *YPK2*-2X FLAG into the *natMX4* vector, the *YPK2* promoter, *YPK2* coding sequence without the STOP codon and the terminator were amplified from the genomic DNA. The forward primer to amplify the *YPK2* terminator was carrying the 2X FLAG sequence with the STOP codon upstream of the sequence that was specific to the *YPK2* terminator. The PCR products were cloned into the empty pBV133 plasmid linearized by *Sac*I and *Spe*I. To clone the *YPK2* allele containing the D265A mutation that was made by analogy with the Tor2-independent *S. cerevisiae* mutation D239A (KAMADA *et al*. 2005) in the *ScYPK2*, the 754-bp fragment of mutated DNA was synthesized by Genscript Biotech Corporation and introduced into the pLS23 that was first linearized by *Mfe*I and *Eag*I. The correct clone was identified based on the elimination of *Mfe*I site in the mutant allele and verified by sequencing. To clone the *YPK2* allele containing the K399A mutation that was made in analogy with the *S. cerevisiae* mutation K376A in *ScYPK1*, the 842-bp fragment of mutated DNA was synthesized by Genscript Biotech Corporation and introduced into the pLS23 that was first linearized by *Avr*II and *Kas*I. The correct clone was identified based on the elimination of the *Avr*II site in the mutant allele and verified by sequencing. We used an artificial reporter gene that has been shown to be regulated by ergosterol levels corresponding to a transcriptional fusion between 5 concatemerized sterol response elements (SREs) from the *ERG1* promoter (referred to as 5X SRE) that we have previously demonstrated is responsive to Upc2A and ergosterol starvation (VU AND MOYE-ROWLEY 2022). To generate the *pUC19* vector containing the *natMX4*-5X SRE-*CYC1*-*lacZ*- integration cassette, all DNA fragments were first PCR amplified – (i) pUC19 backbone of the construct was amplified from the respective pUC19, (ii) *natMX4* coding sequence under the *TEF1* promoter and (iii) *TEF1* terminator were amplified from plasmid pBV133 and (iv) the 5X SRE-*CYC1*-*lacZ* fragment was amplified from pLS27. The identity of the construct was confirmed by restriction analysis using *Not*I and *Sac*II and the linearized construct was integrated into the *HO* locus.

**Table 2.**
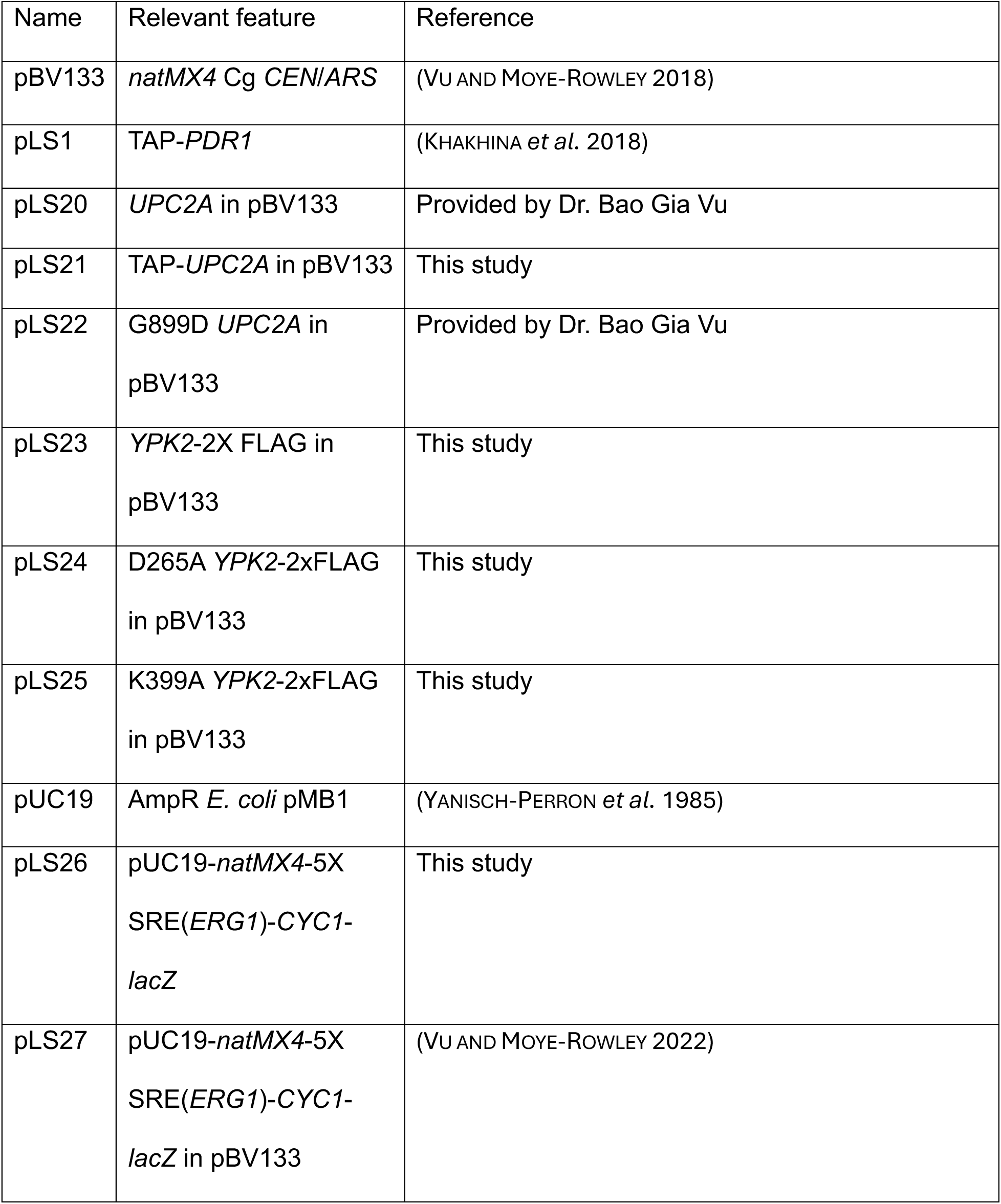
Plasmids used in this work.

| Name | Relevant feature | Reference |
| --- | --- | --- |
| pBV133 | *natMX4* Cg *CEN/ARS* | (VU AND MOYE-ROWLEY 2018) |
| pLS1 | TAP-*PDR1* | (KHAKHINA *et al.* 2018) |
| pLS20 | *UPC2A* in pBV133 | Provided by Dr. Bao Gia Vu |
| pLS21 | TAP-*UPC2A* in pBV133 | This study |
| pLS22 | G899D *UPC2A* in pBV133 | Provided by Dr. Bao Gia Vu |
| pLS23 | *YPK2-2X* FLAG in pBV133 | This study |
| pLS24 | D265A *YPK2-2xFLAG* in pBV133 | This study |
| pLS25 | K399A *YPK2-2xFLAG* in pBV133 | This study |
| pUC19 | AmpR *E. coli* pMB1 | (YANISCH-PERRON *et al.* 1985) |
| pLS26 | pUC19-*natMX4-5X*<br>SRE(*ERG1*)-*CYC1-lacZ* | This study |
| pLS27 | pUC19-*natMX4-5X*<br>SRE(*ERG1*)-*CYC1-lacZ* in pBV133 | (VU AND MOYE-ROWLEY 2022) |

### Transformation of *C. glabrata*

The chemical yeast transformation was performed using the lithium acetate method (GIETZ AND WOODS 2002). 3 OD_600_ units of mid-log cells and 500 ng of plasmid DNA were used per transformation reaction. Cells were exposed to heat shock at 42°C for 1 hour and plated on selective synthetic complete media or incubated in the liquid YPD at 30°C overnight and plated on the YPD media containing nourseothricin the next day. To disrupt genes or integrate DNA cassettes, the CRISPR-Cas9 method of transformation was used as referred in (GRAHL *et al*. 2017). To select transformants, plates were incubated at 30°C for 2 days and individual colonies were PCR-verified for targeted integration of the transformation construct.

### TAP-tag purification and mass spectrometry

The protocol was performed as described in (RIGAUT *et al*. 1999). 600 OD_600_ units of mid-log cells grown in YPD with nourseothricin were harvested and resuspended in 500 μL of NP-40 lysis buffer (15 mM Na_2_HPO_4_, 10 mM NaH_2_PO4, 1% Nonidet, 150 mM NaCl, 2 mM EDTA) complemented by 1 mM PMSF, 1 mM DTT, 1x Complete Protease Inhibitor, 50 mM sodium fluoride, and 0.1 mM sodium vanadate. Cells were lysed in the presence of glass beads (0.5 mm) in 4 cycles of vortexing for 5 min at maximum speed followed by cooling on ice for 1 min. Lysates were cleared by centrifugation, and the supernatant was loaded onto an IgG–sepharose column, followed by incubation at 4°C for 4h. Beads were washed (25 mM Tris-HCl pH 8.0, 0.1% Nonidet, 300 mM NaCl) followed by the second wash using the same buffer containing 150 mM NaCl. TAP-Upc2A proteins were eluted by cleavage with TEV protease (100 U) overnight at 4°C. Eluates were then applied onto the calmodulin–sepharose column in the presence of 2 mM CaCl_2_, incubated for at 4 °C for 2 h followed by washing with the same buffer. Bound Upc2A proteins were eluted from the column by addition of 20 mM EGTA. The eluates were TCA-precipitated, washed with acetone, and dried. Precipitates were solubilized in 50 μl 50 mM triethylammonium bicarbonate (TEAB) in 50:50 (v/v) trifluoroethanol (TFE)/H_2_O. During heat denaturation (90°C, 10 min) the proteins were reduced and alkylated by tris(2-carboxyethyl)phosphine at a final concentration of 10 mM and 2-chloroacetamide at 12.5 mM. Subsequently the samples were evaporated in a vacuum concentrator (Eppendorf). Proteins were resolubilized in 50 μl with 100 mM TEAB in 5:95 (v/v) TFE/H_2_O and digested overnight (18 h, 37°C) with a Trypsin/LysC mixture (Promega) at a protein to protease ratio of 25:1. Tryptic peptides were dried in a vacuum concentrator, resolubilized in 30 μl of 0.05% TFA in H_2_O/acetonitrile 98/2 (v/v) and filtered through spin filters with a hydrophilic PTFE membrane 0.2 µm (Merck Millipore). The filtrate was transferred to HPLC vials and injected into the LC-MS/MS instrument.Raw mass-spectrometry data containing the list of all Upc2A-interacting proteins is provided in supplementary table 1.

### LC-MS/MS analysis

Each sample was measured in triplicate (3 analytical replicates). LC-MS/MS analysis was performed on an Ultimate 3000 nano RSLC system connected to an Orbitrap Exploris 480 mass spectrometer (both Thermo Fisher Scientific, Waltham, MA, USA) with FAIMS. Peptide trapping for 5 min on an Acclaim Pep Map 100 column (2 cm x 75 µm, 3 µm) at 5 µL/min was followed by separation on an analytical Acclaim Pep Map RSLC nano column (50 cm x 75 µm, 2µm). Mobile phase gradient elution of eluent A (0.1% (v/v) formic acid in water) mixed with eluent B (0.1% (v/v) formic acid in 90/10 acetonitrile/water) was performed using the following gradient: 0 min at 4% B, 6 min at 8% B, 30 min at 12% B, 75 min at 30% B, 85 min at 50% B, 90-95 min at 96% B, 95.1-120 min at 4% B. Positively charged ions were generated at a spray voltage of 2.2 kV using a stainless steel emitter attached to the Nanospray Flex Ion Source (Thermo Fisher Scientific). The instrument was operated in data-dependent MS2 mode. Precursor ions were monitored at *m/z* 300-1200 at a resolution of 120,000 FWHM (full width at half maximum) using a maximum injection time (ITmax) of 50 ms and 300% normalized AGC (automatic gain control) target. Precursor ions with a charge state of z=2-5 were filtered at an isolation width of *m/z* 4.0 for further fragmentation at 28% HCD collision energy. MS2 ions were scanned at 15,000 FWHM (ITmax=40 ms, AGC= 200%). 2 different compensation voltages were applied (-50V, -70V).

### Protein database search

Tandem mass spectra were searched against the UniProt database of *Candida glabrata* (https://www.uniprot.org/proteomes/UP000002428; 2023/03/22) using Proteome Discoverer (PD) 3.0 (Thermo) and the database search algorithms Mascot 2.8, Comet, MS Amanda 2.0, Sequest HT with and without INFERYS Rescoring, and CHIMERYS. Two missed cleavages were allowed for the tryptic digestion. The precursor mass tolerance was set to 10 ppm and the fragment mass tolerance was set to 0.02 Da. Modifications were defined as dynamic Met oxidation, phosphorylation of Ser, Thr, and Tyr, acetylation of Ser, Thr, Tyr, Arg and Lys as well as static Cys carbamidomethylation. A strict false discovery rate (FDR) < 1% (peptide and protein level) was required for positive protein hits. Furthermore, search engine scores were filtered for either Mascot (>30), Comet (>3), MS Amanda 2.0 (>300), Sequest HT (>3), or CHIMERYS (>2). The Percolator node of PD3.0 and a reverse decoy database was used for q-value validation of spectral matches. Only rank 1 proteins and peptides of the top scored proteins were counted. Label-free protein quantification was based on the Minora algorithm of PD3.0 using the precursor abundance based on intensity and a signal-to-noise ratio >5.

### Data Availability

The mass spectrometry proteomics data have been deposited to the ProteomeXchange Consortium via the PRIDE partner repository (PEREZ-RIVEROL *et al*. 2025) with the dataset identifier PXD078067 and 10.6019/PXD078067.

### Drug treatment

Mid-log cells were spotted in 10-fold serial dilutions on YPD containing the following drugs: fluconazole (LKT laboratories), myriocin (0.25 μg/mL) (Sigma-Aldrich), caspofungin (0.05 μg/mL) (APExBIO), amphotericin B (1 μg/mL) (Sigma-Aldrich). Cells treated with fluconazole in the liquid YPD media were exposed to the final concentration of 20 μg/mL at 30°C for 2 hours.

### Real time qPCR

Five OD_600_ units of mid-log cells were used per sample. Total RNA was extracted using Trizol (Invitrogen) and chloroform. RNA was purified with RNeasy Mini Kit (Qiagen) and 500 nanograms of total RNA were used to make cDNA using iScript cDNA Synthesis kit (Bio-Rad). qPCR was performed using iTaq Universal SYBR Green Supermix (Bio-Rad). The average Ct value for each sample was calculated from the triplicate. 18S rRNA was used for normalization of variable cDNA levels. All samples were further normalized to untreated wild type strain. A comparative 2^−ΔΔCt^ method was used to calculate the fold change of the gene of interest between samples (LIVAK AND SCHMITTGEN 2001). All measurements represent the result of two independent experiments and the error bars were calculated as standard error of the mean.

### Western blot analysis

3 OD_600_ units of mid-log cells were used per sample. Proteins were extracted as previously described (SHAHI *et al*. 2010), resuspended in urea sample buffer (8 M urea, 1% 2-mercaptoethanol, 40 mM Tris-HCl pH 8.0, 5% SDS, bromophenol blue) and incubated at 37°C for 1 hour. The resuspended proteins were boiled at 95°C for 10 minutes and aliquots were resolved on precast Express Plus 4–12% Gradient Gel (GenScript). Proteins were transferred to nitrocellulose membrane, blocked with 5% nonfat dry milk and probed with primary antibody. All membranes were also probed for tubulin with 12G10 anti-alpha-tubulin monoclonal antibody

(Developmental Studies Hybridoma Bank at the University of Iowa) as a normalization control. Secondary Li-Cor antibodies IRD dye 680RD goat anti-rabbit and IRD dye 800LT goat anti-mouse were used in combination with the Li-Cor Infrared Imaging System and Image Studio Lite software (Li-Cor) to detect and quantify the signal from the western blot. The relative protein levels were normalized to tubulin levels of the corresponding strain and then compared to the reference strain.

### Chromatin immunoprecipitation and qRT-PCR

The chromatin immunoprecipitation (ChIP) experiment was performed as previously described (VU *et al*. 2019). 50 OD_600_ units of mid-log cells were treated with 1% formaldehyde and the reaction was inhibited by glycine. Cells were resuspended in FA-lysis buffer (50 mM HEPES-KOH, 140 mM NaCl, 1 mM EDTA, 1% Triton X-100, 0.1% sodium deoxycholate, supplemented with 1 mM PMSF and 1x Complete protease inhibitor). The cell suspension was vortexed with glass beads (0.5 mm) at 4°C for 20 minutes. The beads were separated by centrifugation, the lysate and pellets were resuspended and 2×250 ml volumes were aliquoted into 1.5 mL Bioruptor Pico Microtubes and sheared by Diagenode Bioruptor Pico instrument (Diagenode) using the following conditions: sonication ON for 30 sec, sonication OFF for 30 Sec, number of cycles 15, total running time 15 mins/per 6 samples. Next, the supernatant was separated from the rest of the sample by centrifugation and an aliquot was removed as input control. For immunoprecipitation, anti-Upc2A polyclonal antibody was used (1:100). The antibody was first preincubated with the lysate at 4°C for 2 hours. Next, Dynabeads Protein A Magnetic Beads (Invitrogen) were added to the sample and incubated at 4°C overnight on a nutator. Washing and all subsequent steps were performed as described previously (CHUNG *et al*. 2014). To perform qRT-PCR on the purified DNA, primers specific to *ERG11* promoter (-561 to -694 relative to start codon), and *ERG1* promoter (-615 to -916 relative to start codon) were used. Each sample was analyzed in triplicate. 1 μl of ChIP-ed DNA and 10-fold diluted input DNA was used in the reaction with the final volume of 20 μl. The PCR reaction was carried out under the following conditions: One cycle of 95°C for 30 seconds, 40 cycles of 95°C for 15 seconds and 56°C for 30 seconds on MyiQ2 BioRad instrument. To calculate the signal of enriched DNA of the respective promoter, the percent input method was applied. The data represent the result of two independent experiments and the error bars were calculated as standard error of the mean.

### Beta-galactosidase Assay

Cells grown at 30°C overnight in YPD media were back-diluted in the morning and grown to a mid-log phase of OD_600_ = ∼ 1. Next, 1 mL of culture was spun down and pellet was resuspended in 1 mL of Z-buffer (60 mM Na_2_HPO_4_, 40 mM NaH_2_PO_4_, 10 mM KCl, 1 mM MgSO_4_, 50 mM 2-Mercaptoethanol) + 20 µl 0.1% SDS. As a control sample, 1 ml of Z-buffer with no cells was used. Afterwards, 30 μl of chloroform was added and the samples were vortexed at high speed for 10 seconds. Before the assay, tubes were warmed at 30°C for 3 minutes. Next, 200 μl of 4 mg/mL o-nitrophenyl-beta-D-galactopyranoside (ONPG) in ddH20 was added and the colorimetric reaction was stopped after 10 minutes of incubation at 30°C by adding 500 μl of 1M Na_2_CO_3_. The cells were spun down and the absorbance of the supernatant was assessed at A_420_ to measure beta-galactosidase activity. Each measurement was normalized to A_600_ of the respective strain and the specific activity was calculated as Miller units based on the equation: 1000 x A_420_ / volume of cells (1 mL) x A_600_ x time (10 min). The data represent the result of two independent experiments and the error bars were calculated as standard error of the mean.

### RNA-sequencing

We prepared total RNA from YPD-grown cells treated with antifungal drugs as described previously (CONWAY *et al*. 2024) . Briefly, wild-type and *ypk2*Δ cells were grown to mid-log phase and split between three culture conditions for varying lengths of time: (1) 16 µg/ml fluconazole for three hours, (2) 0.25 µg/ml amphotericin B for one hour, and (3) 50 ng/ml caspofungin for one hour. Six OD_600_ units of cells were pelleted from each pre- and post-treatment culture, fixed in Trizol (Invitrogen), and total RNA was purified using a RNeasy Mini Kit (Qiagen). RNAseq was performed by Plasmidsaurus (Louisville, KY) using Illumina Sequencing Technology and a 3’ end counting approach for analysis of transcript identity and abundance. All conditions were assayed in biological triplicate, and the results of the three independent experiments were averaged for each condition prior to comparative analyses. These data will be submitted to the GEO database.

### Statistics

Statistical significance was assessed by unpaired Student *t*-test. Significance in each comparison was indicated as follows: \**P* < 0.05, \*\**P* < 0.01, and \*\*\**P* < 0.001.

## Results

### Co-purification of protein kinase Ypk2 with Upc2A

While the induction of Upc2A-dependent gene expression is well-established, the mechanism(s) underlying this increase in activity are not known. To gain insight into the proteins that associate with Upc2A and may act to modulate its function, we expressed an amino-terminally fused tandem affinity purification (TAP)-Upc2A and validated that this chimeric protein retained normal function (Supplementary Figure 1). Protein extracts were prepared from *upc2AΔ* cells containing this TAP-*UPC2A* fusion gene after growth with or without the presence of fluconazole. After TAP purification, mass spectrometry was performed on the purified proteins to identify co-purifying polypeptides. We detected a number of proteins that co-purified with TAP-Upc2A (STab_Proteomics.xlsx) but will focus here on the characterization of the AGC protein kinase Ypk2.

To confirm our co-purification results, we generated a 2X FLAG-tagged allele of *YPK2* and introduced this into cells containing an HA-tagged allele of *UPC2A* or a *upc2AΔ* strain. Appropriate transformants were grown to mid-log phase and then whole cell protein extracts produced under non-denaturing conditions. These lysates were then subjected to immunoprecipitation using anti-HA antibodies. The resulting immunoprecipitates were then analyzed using SDS polyacrylamide gel electrophoresis, followed by western blotting using either anti-HA or anti-FLAG antibodies. Immunoreactive polypeptides were detected with corresponding secondary antibodies (Figure 1A).

**Figure 1.**
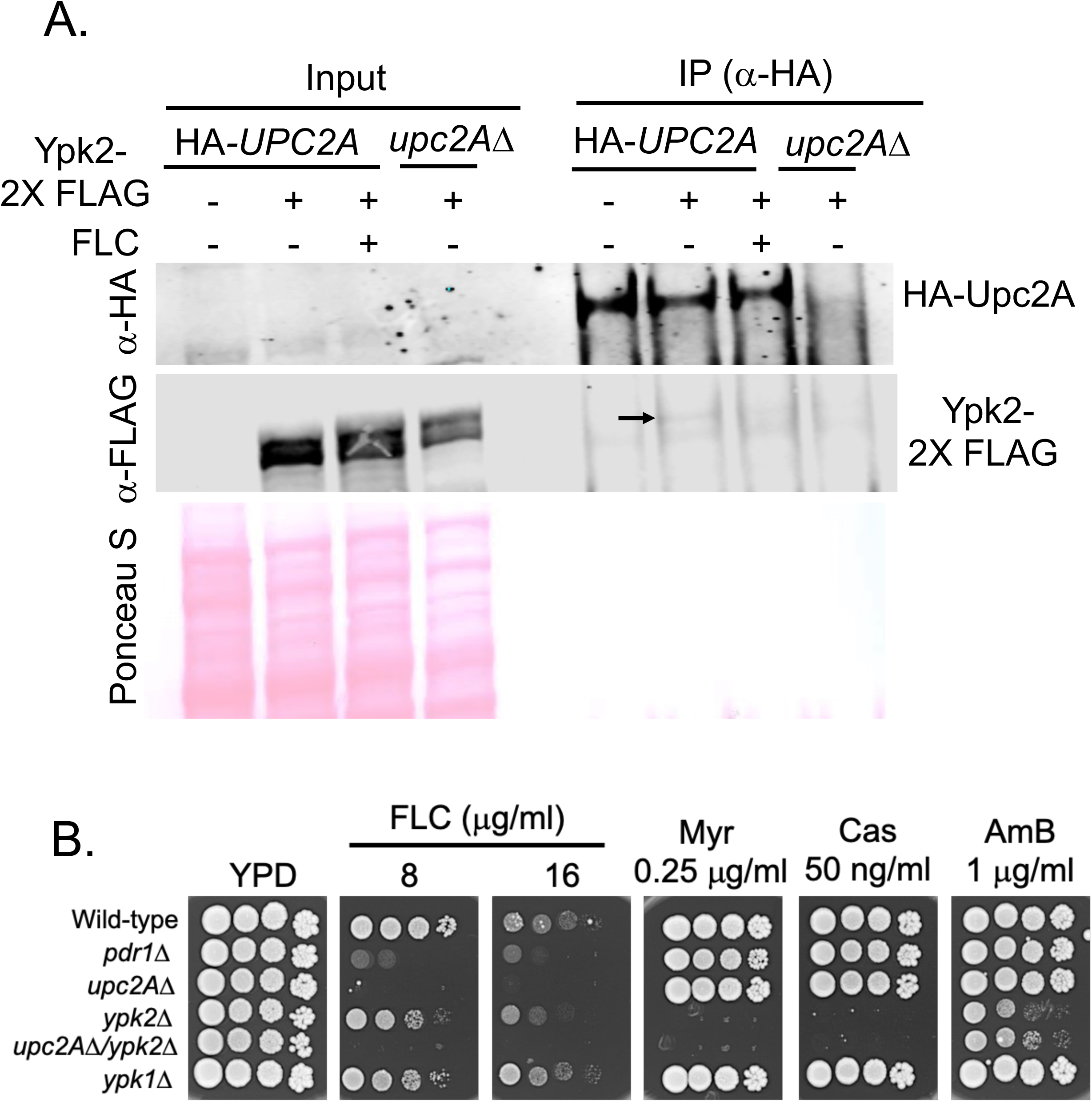
Protein kinase Ypk2 that co-purifies with Upc2A is important for fluconazole resistance. A. Western blot following the co-immunoprecipitation confirming that Upc2A (N-terminally HA-tagged) copurifies with Ypk2 (C-terminally 2x FLAG tagged). Input corresponds to the Upc2A or Ypk2 present in the 1% fraction of the whole cell lysate. The arrow indicates to the band corresponding to the Ypk2-2X Flag protein that co-purified with HA-Upc2A during the pulldown using the α-HA antibody. Ponceau staining of the western blot membrane illustrates the equal loading of the whole cell lysates among all four tested strains. B. Differential drug susceptibility profiles of key *C. glabrata* strains. Isogenic derivatives of the wild-type *C. glabrata* strain KKY2001 were grown to mid-log phase and then assayed by serial dilutions on rich media (YPD) containing the indicated concentrations of different drugs. Abbreviations are FLC, fluconazole; Myr, myriocin; Cas, caspofungin and AmB, amphotericin B. Plates were incubated for two days at 30°C and then photographed.

Low but detectable levels of Ypk2-FLAG were found in HA-Upc2A immunoprecipitates supporting the conclusion that Ypk2 and Upc2A associate in the cell. Difficulty in recovering kinase-substrate partners is well-documented (reviewed in (DE OLIVEIRA *et al*. 2016; SUGIYAMA *et al*. 2019)) as these interactions are transient. This is likely to explain the low levels of co-immunoprecipitated Ypk2 found in the Upc2A immunoprecipitates. In these experiments, we did not find any indication that the presence of the antifungal drug fluconazole modulated the degree of interaction between Ypk2 and Upc2A.

To extend this observation, we wanted to evaluate the antifungal drug phenotypes of cells lacking *YPK2* and its homologue *YPK1*. Isogenic strains were generated that lacked either of these protein kinase-encoding genes as well as *PDR1* or *UPC2A*. These strains were tested on fluconazole, caspofungin, myriocin and amphotericin B. All strains were grown to mid-log and then serial dilutions of each strain placed on rich YPD medium plates containing the indicated concentrations of each drug (Figure 1B).

Loss of *YPK2* caused an increase in fluconazole, caspofungin, myriocin and amphotericin B susceptibility while loss of *YPK1* only increased fluconazole susceptibility. Loss of *UPC2A* increased fluconazole susceptibility while a double *upc2AΔ ypk2Δ* null did not show any additional sensitivities compared to the single *ypk2Δ* strain. The *pdr1Δ* strain was highly susceptible to fluconazole but had no additional sensitivities under these conditions.

Given that loss of Ypk2 impacted fluconazole resistance, we examined the fluconazole-dependent induction of downstream gene expression in isogenic wild-type and *ypk2Δ* strains. Strains were grown in YPD or challenged with the addition of fluconazole. After 3 hours of growth, cells were harvested and total RNA prepared. Using appropriate primers, levels of the indicated transcripts were assessed by RT-qPCR.

We first measured RNA levels for *ERG11* and *CDR1*, two well-established genes that impact azole susceptibility (Figure 2A). Loss of Ypk2 prevented fluconazole induction of *ERG11* transcription while this had no significant effect on *CDR1* drug induction. We then examined fluconazole induction of two *ERG* genes that act earlier in the ergosterol biosynthetic pathway (*ERG10*, *ERG1*) as well as later (*ERG2*) relative to *ERG11*. All three of these genes were induced upon fluconazole treatment (Figure 2B). While induction of both *ERG1* and *ERG10* were reduced in *ypk2Δ* cells, *ERG2* induction was not significantly affected. These data support the view that fluconazole-induced Upc2A gene activation is blunted in the absence of Ypk2 and that *ERG* gene regulation involves factors beyond Upc2A as reported earlier (OLLINGER *et al*. 2021).

**Figure 2.**
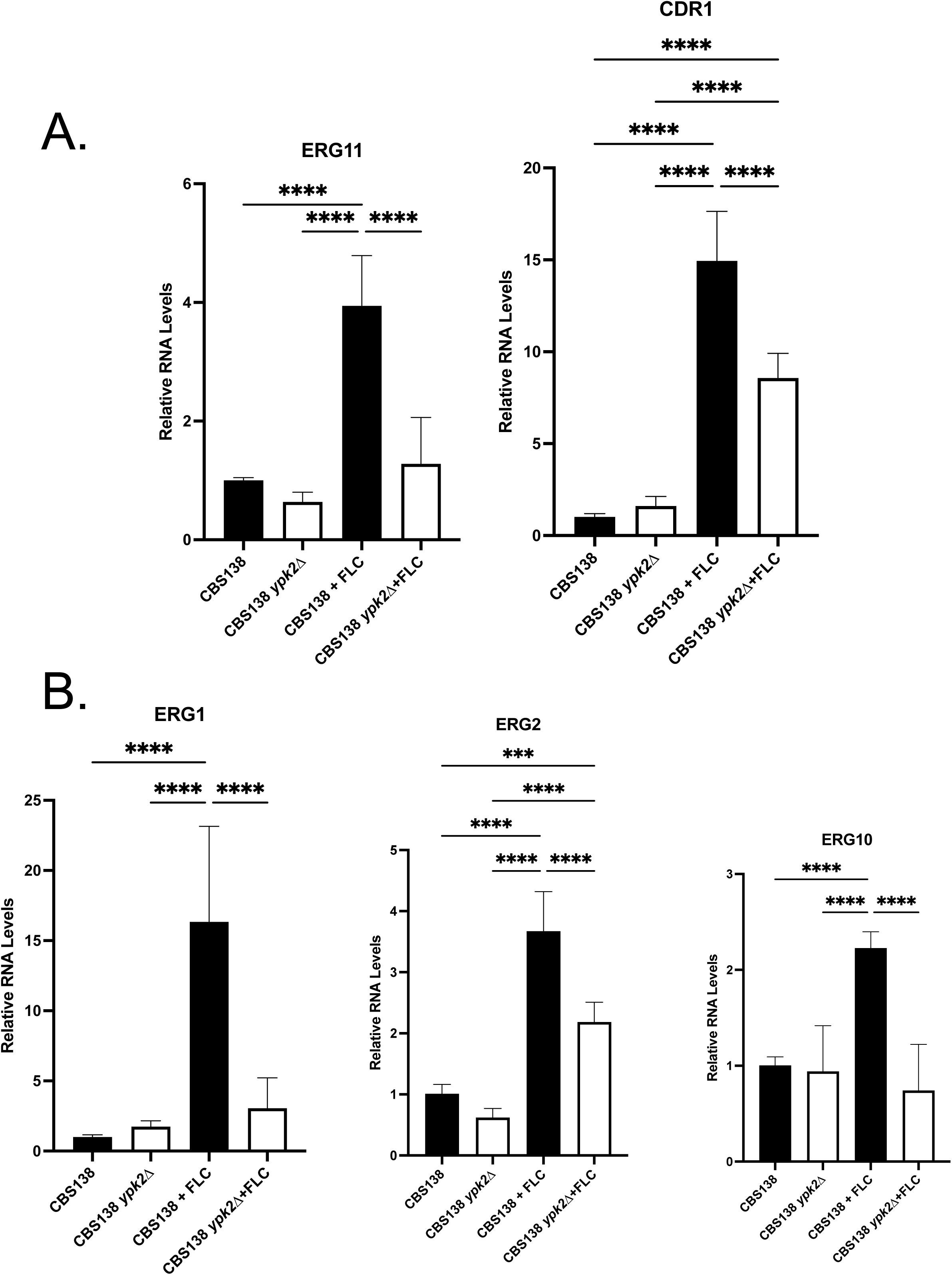
Loss of Ypk2 inhibits fluconazole-induced expression of *ERG* genes. Wild-type and *ypk2Δ* cells were grown with or without fluconazole (20 μg/ml for two hours). Total RNA was then prepared and analyzed by RT-qPCR using primer pairs that detect the indicated genes. A. Expression of *ERG11* was induced in the presence of fluconazole and this induction was significantly reduced upon loss of Ypk2. Fluconazole induction of *CDR1* was not significantly impacted by loss of Ypk2. B: Fluconazole induction of *ERG1* and *ERG2* (both members of the late ergosterol biosynthetic pathway, as is *ERG11*) was assayed for RT-qPCR as was transcription of *ERG10* (encodes the first committed step in ergosterol biosynthesis) to compare their requirement for Ypk2.

### Evidence for Ypk2 activation of Upc2A function

To assess the mechanism underlying Ypk2 regulation of Upc2A activity, we examined several properties of this transcription factor. We analyzed DNA-binding of Upc2A by carrying out chromatin immunoprecipitation (ChIP) in isogenic wild-type and *ypk2Δ* cells, grown under control of fluconazole-challenged conditions. Fixed chromatin prepared under these growth conditions was sheared, DNA bound to Upc2A recovered by immunoprecipitation using anti-Upc2A antibodies (VU *et al*. 2021) and the crosslinks reversed. Levels of the *ERG11* promoter were assessed using qPCR and appropriate primers (Figure 3A).

**Figure 3.**
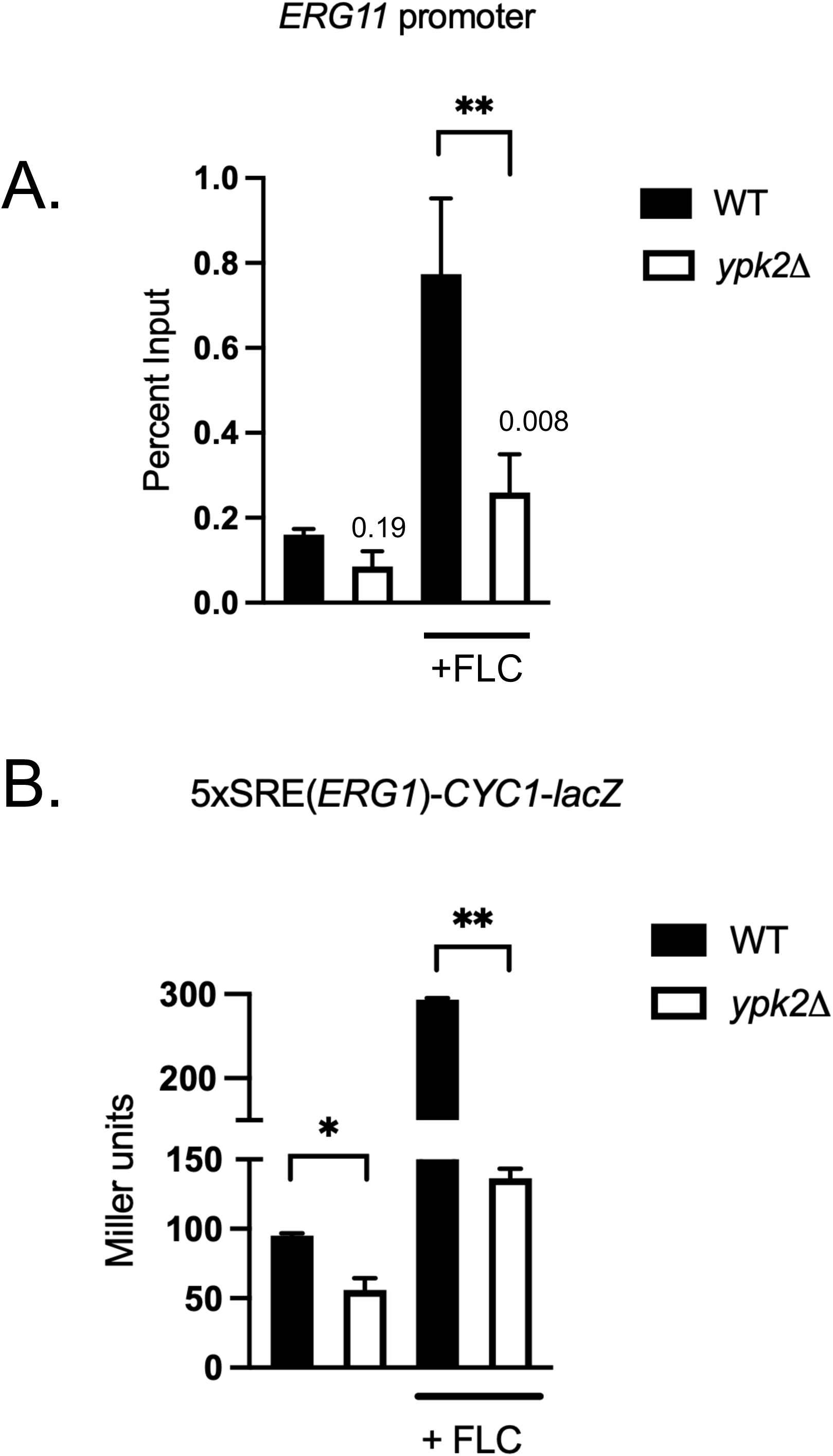
Upc2A binding to *ERG11* promoter and transcriptional activation is reduced upon loss of Ypk2. A. Chromatin immunoprecipitation assay comparing the binding of Upc2A to the *ERG11* promoter in response to the loss of Ypk2. Fixed chromatin was prepared from the indicated strains in the presence or absence of fluconazole, Percent input represents the amount of DNA from the *ERG11* promoter recovered by the α-Upc2A antibody in the ChIP reaction relative to total *ERG11* promoter DNA present in the fractions. B. An 5X sterol response element (SRE) from the *ERG1* promoter concatemerized and placed upstream of the Sc *CYC1* promoter fused to *E. coli lacZ* was integrated into the HO locus of wild-type and isogenic *ypk2Δ* cells. Appropriate transformants were grown in the presence or absence of fluconazole. Levels of b-galactosidase activity produced in these strains were determined by a standard assay (presented as Miller units on the ordinant).

As we have seen before (VU *et al*. 2021), the level of *ERG11*-bound Upc2A was enhanced upon fluconazole treatment in wild-type cells. However, when this analysis was performed using *ypk2Δ* -derived chromatin, the fluconazole-induced level of Upc2A-bound *ERG11* promoter DNA recovered was decreased by 4-fold. These data indicate that Ypk2 function is required for the normal increase in Upc2A DNA-binding seen upon fluconazole treatment.

Next, we prepared an artificial *lacZ* reporter in which 5 copies of the Upc2A binding site from the *ERG1* gene (Sterol response element: SRE) were concatemerized and placed upstream of a *Saccharomyces cerevisiae CYC1* proximal promoter region that was translationally fused to *E. coli lacZ*. This construct was integrated into either wild-type or *ypk2Δ* cells. Appropriate transformants were grown in the presence or absence of fluconazole and *CYC1*-dependent levels of *lacZ*-encoded β-galactosidase measured (Figure 3B).

Levels of β-galactosidase were induced upon fluconazole treatment by 3-fold in wild-cells from 100 to nearly 300 Miller units. When these same assays were done in a *ypk2Δ* strain, the SRE-dependent β-galactosidase activities varied from 50 Miller units to 140 Miller units in the presence of fluconazole. We have previously shown that this fusion gene serves as a sensitive reporter for the levels of Upc2A-dependent transactivation (VU AND MOYE-ROWLEY 2022). These data are consistent with Upc2A-dependent transcriptional activation being reduced in the absence of Ypk2 activity.

Together, these data suggest that the activity of Upc2A is increased by Ypk2 function with some of this increase likely being at the level of DNA binding.

### Loss of Ypk2 can be largely suppressed by a hyperactive *UPC2A* allele

We have previously described the G899D allele of *UPC2A* that exhibits constitutively elevated Upc2A function. This mutant was developed based on a previously described mutation isolated in the *Saccharomyces cerevisiae* homologue *UPC2* (CROWLEY *et al*. 1998) that led to this factor driving elevated target gene expression which we found was also true for *C. glabrata* G899A Upc2A (VU *et al*. 2021). To evaluate the effect of loss of Ypk2 function on G899A Upc2A, we prepared isogenic *upc2AΔ* and *upc2AΔ ypk2Δ* strains. These strains were transformed with low-copy-number plasmids expressing either the wild-type or G899D forms of Upc2A or the vector only. Transformants were tested for their ability to grow in the presence of fluconazole (Figure 4A).

**Fig. 4.**
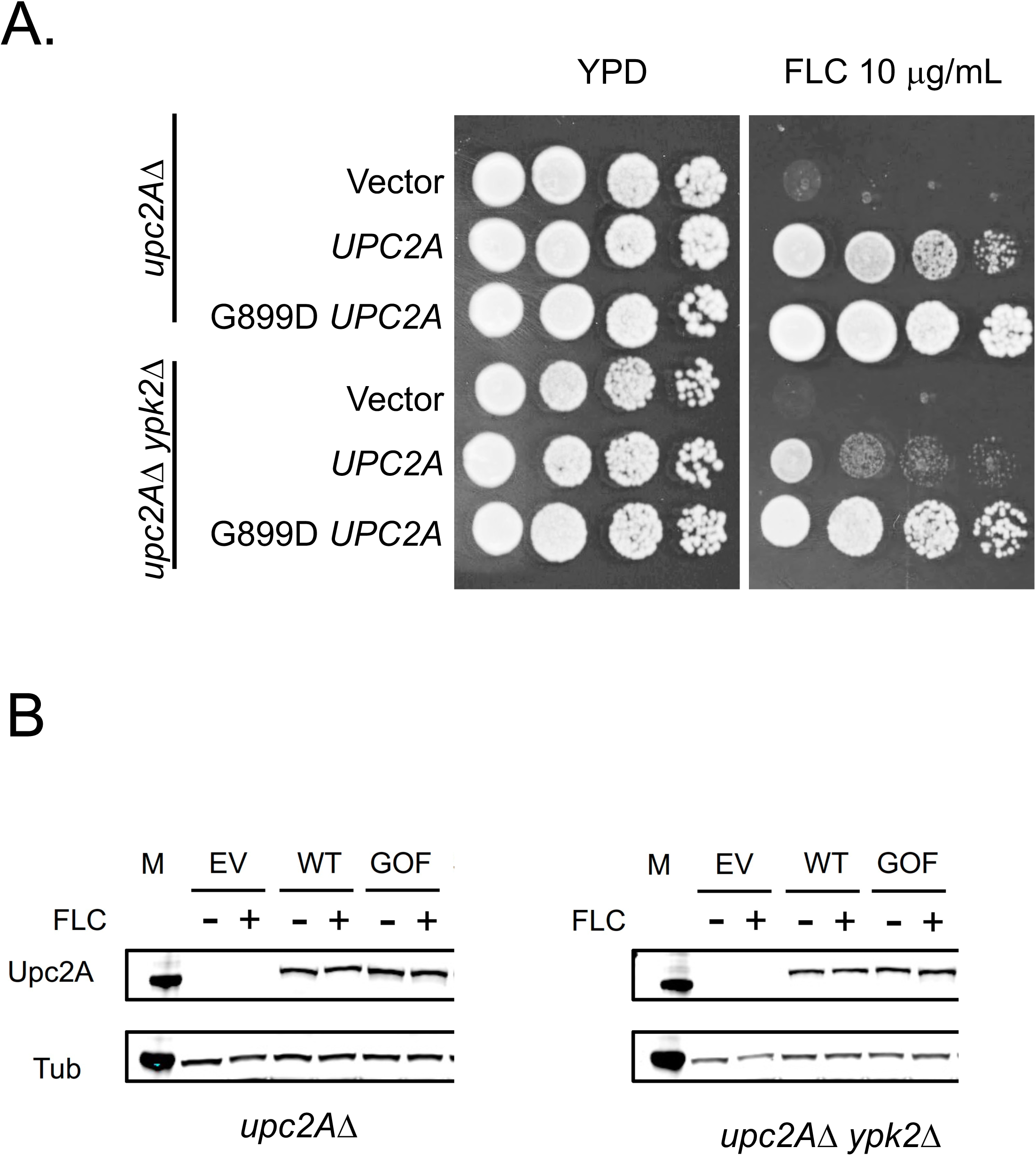
Hyperactive *UPC2A*-G899D mutant is less affected by the loss of *YPK2* than wild-type. A. Isogenic *upc2A*Δ and *upc2AΔ ypk2Δ* strains were transformed with low-copy-number plasmids containing wild-type or a gain-of-function form (G899D) of *UPC2A* as well as the empty vector plasmid (Vector). Transformants were grown to mid-log phase and then assayed by serial dilution plating on rich medium (YPD) or YPD containing 10 μg/ml fluconazole (FLC). Plates were incubated at 30°C and photographed after 48 hours. B. Whole cell protein extracts were prepared from the transformants above and analyzed by western blotting using a a-Upc2A polyclonal antiserum. The absence or presence of fluconazole in the cultures prior to protein extraction is indicated as (-) or (+), respectively. Transformants are abbreviated as EV (Empty Vector), WT (*UPC2A*) or GOF (G899D *UPC2A*).

Loss of Ypk2 strongly increased fluconazole susceptibility when wild-type *UPC2A* was present but had much less impact on the elevated fluconazole resistance displayed by the G899D *UPC2A* transformants. These data suggest that the presence of the G899D *UPC2A* allele relieved much of the reliance of Upc2A function on the presence of Ypk2.

To ensure that the expression levels of these different Upc2A derivatives, in the presence and absence of fluconazole, were comparable, we analyzed the steady-state protein levels of wild-type and G899D Upc2A. Whole cell protein extracts were prepared from the strains listed above, grown with or without fluconazole treatment, and resolved on SDS-PAGE, followed by western blotting with anti-Upc2A antibody (Figure 4B).

Irrespective of the form of Upc2A, the presence of either fluconazole or Ypk2, we did not find any significant change in the steady-state level of Upc2A. These data argue that changes in fluconazole susceptibility are due to a change in the activity state of Upc2A rather than expression.

### Ypk2 kinase activity and regulatory mutations impact phenotypes in *C. glabrata*

We based two different mutant forms of Ypk2 on the extensive literature describing function of the *S. cerevisiae* homologues (reviewed in (JACINTO AND LORBERG 2008; EMMERSTORFER-AUGUSTIN AND THORNER 2023)). Alignments of the sequences of the *C. glabrata* Ypk2 with its *S. cerevisiae* counterpart indicated that the catalytic lysine K399 was the equivalent of the *S. cerevisiae* K373 (ROELANTS *et al*. 2002). This lysine is required for normal catalytic activity of the kinase enzyme. Similarly, the *C. glabrata* aspartate at position 265 is the equivalent of the *S. cerevisiae* Ypk2 D242. Mutation of this amino acid from aspartate to alanine was shown in *S. cerevisiae* to relieve the requirement for TORC2 activation of Ypk2 activity (KAMADA *et al*. 2005). To determine if these positions in the *C. glabrata* kinase had similar roles to those in the *S. cerevisiae* enzyme, we constructed D265A and K399A Cg *YPK2* based on the analogous mutations found in the *S. cerevisiae* gene. These mutant forms of Ypk2 were expressed from low-copy-number plasmids as FLAG-tagged alleles and were transformed into a *ypk2Δ* strain along with the empty vector and the wild-type allele as controls. Appropriate transformants of each type were used for subsequent experiments.

We first tested the ability of each *YPK2* allele to complement the antifungal drug susceptibility phenotypes of the *ypk2Δ* strain (Figure 5A). Both the wild-type and D265A forms of Ypk2 restored normal fluconazole and myriocin susceptibility. The D265A form of Ypk2 exhibited decreased caspofungin susceptibility compared to the wild-type enzyme, suggesting that the activation by TORC2 may limit function of this kinase in the presence of this echinocandin.

**Fig. 5.**
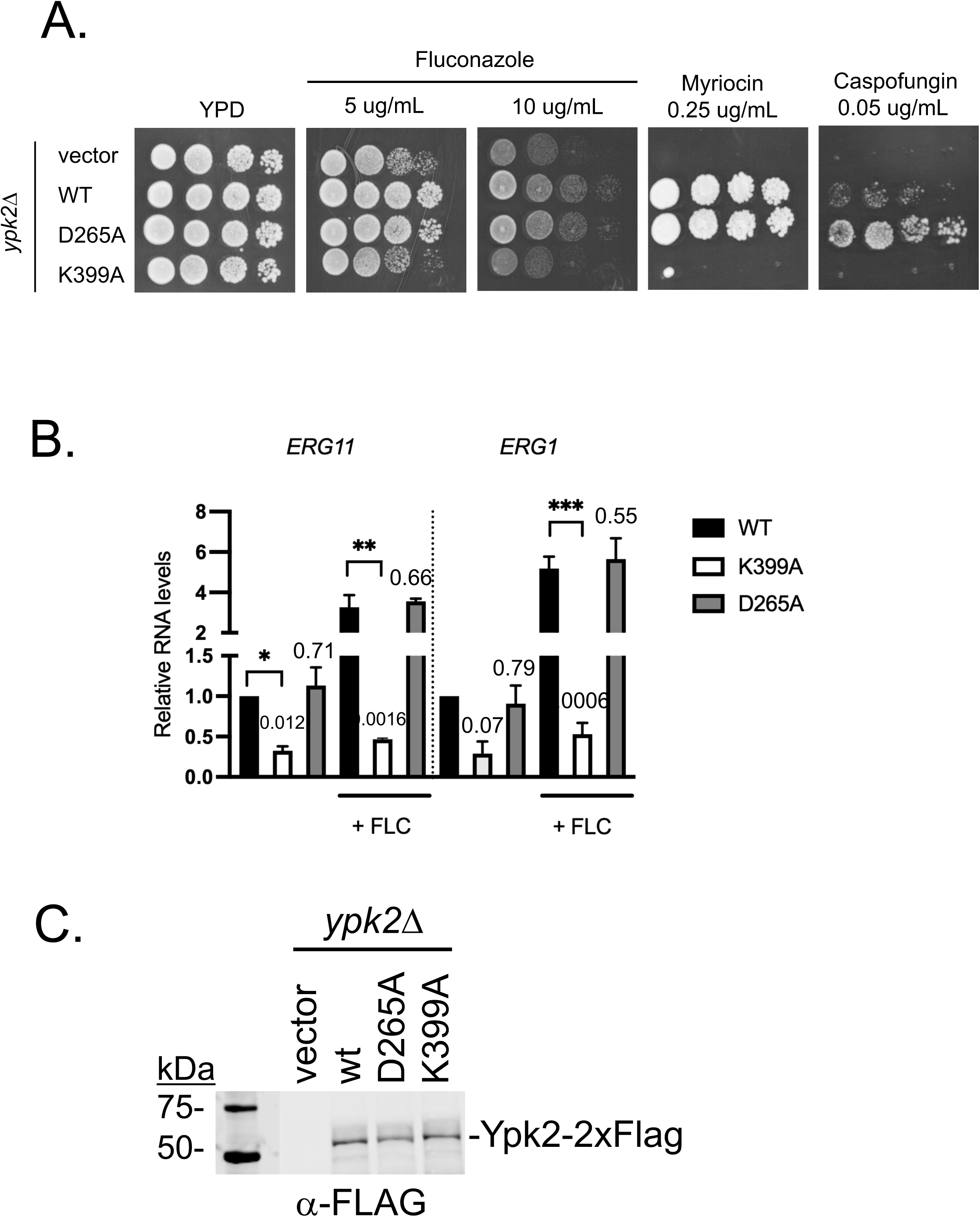
Catalytic activity of Ypk2 is important for fluconazole resistance. A ypk2Δ strain was transformed with a low-copy-number vector (Vector) plasmid or this same plasmid containing the wild-type (wt), D265A or K399A forms of Ypk2. These transformants were used for all experiments here. A. Susceptibility of the Torc2 independent (D265A) and catalytically inactive (K399A) *YPK2* mutants to fluconazole, myriocin and caspofungin were compared to wild-type and empty vector controls by serial dilution on rich medium containing the indicated concentrations of drugs. B. Expression of *ERG1* and *ERG11* in all transformants was determined in the presence or absence of fluconazole. C. Steady-state levels of Ypk2 forms were assessed by western blotting using a mouse monoclonal antibody directed against the C-terminal FLAG tags.

We also examined the regulation of *ERG1* and *ERG11* in the presence of these forms of Ypk2. Transformants from above were grown to mid-log phase, total RNA prepared from untreated or fluconazole-challenged cells and levels of *ERG1* and *ERG11* assayed by RT-qPCR as before (Figure 5B). Both wild-type and D265A forms of Ypk2 normally induced transcription but the K399A kinase defective mutant behaved indistinguishably from the empty vector control.

To ensure that any difference in phenotype from these Ypk2 mutant derivatives was due to a change in the activity of the resulting protein, we analyzed expression of all proteins by western blotting using anti-FLAG antibodies. Transformants were grown to mid-log phase and then equal amounts of whole cell protein extracts resolved on SDS-PAGE followed by transfer to a nylon membrane. These membranes were probed with mouse anti-FLAG and bound antibody visualized by standard technique (Figure 5C).

All these Ypk2 derivatives were expressed at comparable levels, indicating that any phenotypic differences were likely due to changed activity of the mutant protein.

### Transcriptional outputs of Ypk2 vary by stress treatment

Having provided evidence that Ypk2 functions to regulate transcriptional activation by Upc2A with a resulting impact on fluconazole susceptibility, we wanted to determine the Ypk2-regulated transcriptome when cells were challenged with three different stresses. We demonstrated above that cells lacking *YPK2* exhibited increased susceptibility to fluconazole as well as caspofungin and amphotericin B. To determine the similarity of the Ypk2-dependent transcriptomes influenced by these three different antifungal drugs, we carried out RNA-seq analyses of both wild-type and *ypk2Δ* strains treated with each of these drugs. Cells were challenged with each drug under conditions we have used before (CONWAY *et al*. 2024) and total RNA prepared. This RNA was used for RNA-seq analysis and reads mapped back to the *C. glabrata* CBS138 genome using standard approaches.

To assess the effect of each drug treatment, we compared the number of genes that are significantly induced (P-value<0.05) by at least two-fold. The transcriptomic response of these three antifungal genes was comparable in terms of the numbers of genes induced (Figure 6A and Supplementary Table 2). AmB-treated cells exhibited 828 induced genes, while caspofungin- and fluconazole-challenged cells contained 743 and 606 induced genes, respectively. 197 genes were found to be induced in wild-type cells, irrespective of the drug used (This category is labeled Common in Figure 6B). The majority of these commonly induced genes were involved in metabolism and likely reflect the generalized stress caused by these antifungal agents.

**Figure 6.**
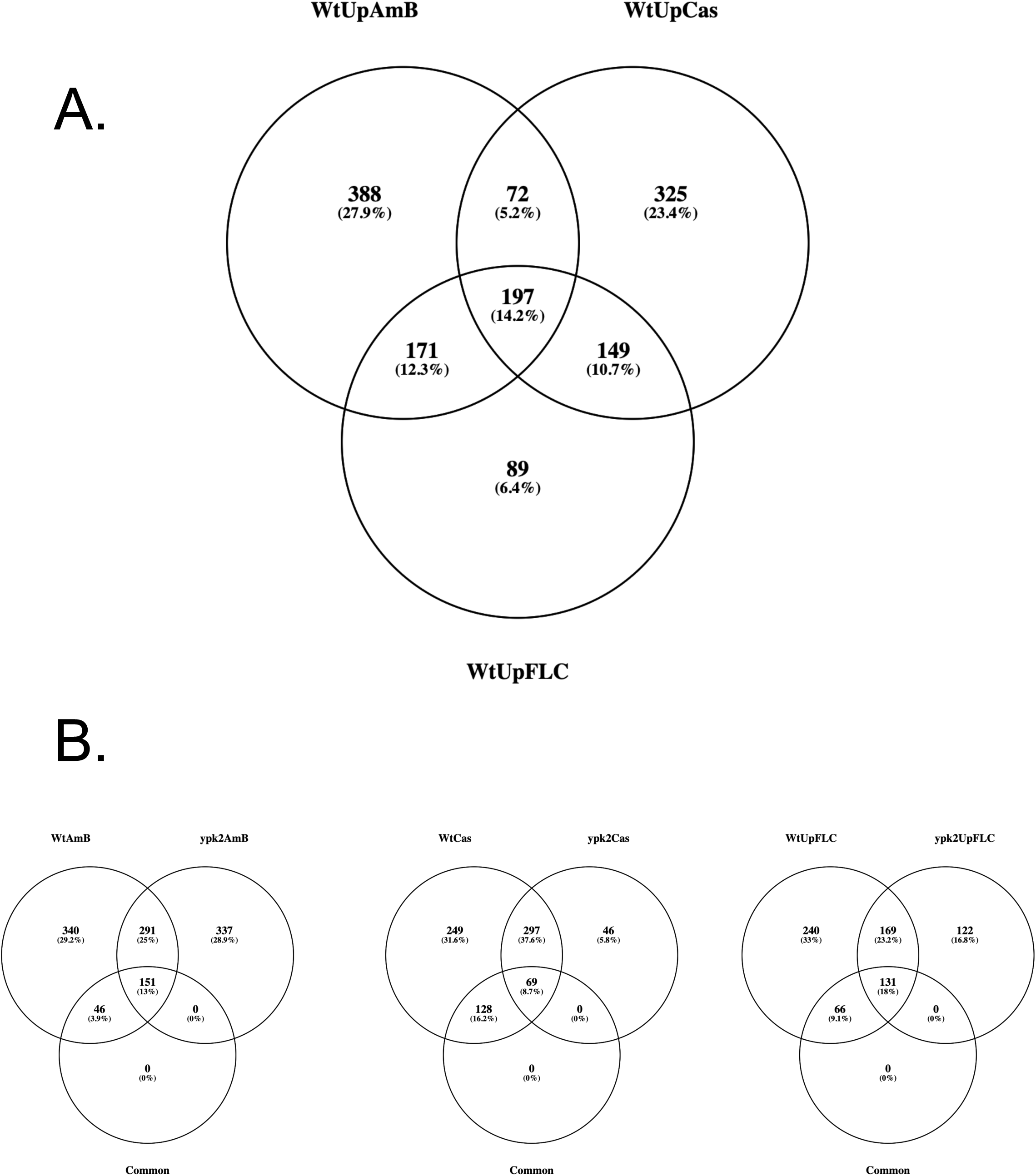
RNA-seq analysis of genes responsive to antifungal drugs and Ypk2. A. Venn diagrams comparing the genes that are induced in wild-type cells by at least two-fold and with a P-value<0.05 when treated with amphotericin B (WtUpAmB), caspofungin (WtUpCas) or fluconazole (WtUpFLC). B. Ypk2-dependent induction of gene expression in response to different antifungal drugs. Genes induced in wild-type cells are from A while those that are induced in *ypk2Δ* cells are indicated. Common genes represent genes that are induced in response to all three antifungal drugs in wild-type cells.

Down-regulated genes that responded to each of these drugs were dominated by repression of protein synthetic apparatus. Translation is well-established to be inhibited during stress (LI *et al*. 1999; SHORE *et al*. 2021), owing to limited energy availability during this type of an environmental challenge. This finding also confirms that our selection of treatment times and amounts of these different antifungal drugs generated a similar general translational inhibition and could be compared in terms of this synthetic block.

We next limited consideration to genes that failed to induce in the *ypk2Δ* background in response to these drug treatments (Figure 6B). We focused on the drug-induced genes that require Ypk2 for their induction as these genes are likely to be involved in the Ypk2-mediated response to the various drug challenges. There were 386 Ypk2-dependent genes in AmB-treated cells, 377 genes in caspofungin-treated cells and 306 in fluconazole-challenged cells. These data indicate the range of the Ypk2-dependent response to these drugs ranges from 300 to 400 genes, approximately. Within the commonly induced genes, the dependence on Ypk2 for induction varies from 46 genes (AmB) to 128 genes (Cas) while 66 genes depend on Ypk2 for induction in the presence of FLC. This behavior shows that, while these common genes are all induced in the presence of these drugs, their mechanism of induction is shaped by each drug through the altered dependence on Ypk2.

We also inspected the Ypk2-dependent, drug-induced genes for shared GO terms to gain insight into the biological processes that were likely changing by altering gene expression. The simplest class of GO term enriched genes was found for caspofungin-induced genes with only 4 categories and the top ranking GO category being carbohydrate metabolic process (Supplementary Table 3). This includes several interesting genes including *PFK26* and *PYC1*, two enzymes involved in gluconeogenesis, along with three different genes encoding cell wall remodeling enzymes: *GAS1*, *SGA1* and *MRX18*. Gas1 has previously been implicated in caspofungin susceptibility in *S. cerevisiae* (LESAGE *et al*. 2004). AmB treatment led to enrichment of GO terms corresponding to activation of genes involved in energy generation, transmembrane transport and nitrogen compound biosynthesis. Fluconazole-induced, Ypk2 dependent genes were also enriched for GO terms corresponding to energy generation and metabolism (as seen for AmB) but also contained 9 genes grouped by GO terminology into lipid modification. These included 6 corresponding to peroxisomal gene products (Pox1, Pxa2, Pex11, Tes1, Mdh3, Eci1), consistent with significant peroxisomal response to ergosterol limitation. Ergosterol has an important role in protein localization to the peroxisomal membrane and function of this organelle (BOUKH-VINER *et al*. 2005; CARMICHAEL AND SCHRADER 2022)

The three additional lipid-related genes are two phosphatases that control lipid production (Ymr1, Ysr3) and a protein involved in steroid detoxification that has been found to be upregulated in azole resistant strains (Atf2). Ymr1 is best described as a regulator of PI3P levels and impacts intracellular trafficking while Ysr3 is involved in regulation of ceramide production (KOBAYASHI AND NAGIEC 2003), a key intermediate in sphingolipid biosynthesis. Sphingolipids interact with ergosterol to maintain proper membrane fluidity and function. Atf2 may be participating in detoxification of ergosterol derivatives that accumulate in the presence of FLC.

## Discussion

These data provide a consistent picture of the protein kinase Ypk2 being an important positive regulator of gene activation by Upc2A. In Sc, Ypk1/2 kinases have been extensively implicated in the post-translational control of regulation of sphingolipid biosynthesis, especially through the modulation of the first committed step in production of these important membrane lipids. Less is known of the role of these kinases in transcription in Sc although experiments using an inhibitable form of Ypk1 have compared the global transcriptome produced with and without acute inhibition of this protein kinase (MACE *et al*. 2020). These experiments predict only roughly 50 genes that are dependent on Sc Ypk1 for fluconazole induction. This observation suggests that the role of Ypk2 in the Cg response to fluconazole treatment may play a more significant role than that seen for Sc Ypk1 in that organism.

The clearest illustration of the impact of Ypk2 on the activity of Upc2A comes from the consideration of the artificial reporter gene in which 5 concatemerized SREs from *ERG1* are placed upstream of the Sc *CYC1-lacZ* gene. Based on previous genetic analyses of *upc2AΔ* strains, *ERG* gene expression in particular and fluconazole susceptibility in general has positive regulators beyond Upc2A (OLLINGER *et al*. 2021). The presence of these additional regulators coupled with the fact that ergosterol biosynthetic enzyme-encoding genes other than *ERG11* impact fluconazole susceptibility, complicates the interpretation of *ERG11* expression as a readout for Upc2A function. The artificial reporter gene contains only SRE as its sole upstream activation sequence and provides a simpler means of monitoring the transcriptional activation of Upc2A isolated from influence of other factors. The reduced level of expression of this reporter gene in the absence of Ypk2 argues for that the impact of this kinase is to stimulate Upc2A-dependent gene activation.

Since Ypk2 stimulates Upc2A function, we used a genetic approach to gain insight into the mechanisms underlying how this kinase modulates Upc2A activity. We previously characterized a GOF form of Upc2A that exhibits constitutively elevated nuclear levels of factor (VU *et al*. 2021). Evidence for the increased nuclear levels of the G899D Upc2A came from the increased DNA-binding to nuclear targets seen in the absence of FLC compared to the wild-type factor. Additionally, while *ERG11*-binding by the wild-type Upc2A increased by 7-fold upon FLC treatment, G899D Upc2A DNA-binding was not further increased in the presence of drug. Based on the work of others (YANG *et al*. 2015; TAN *et al*. 2022), we speculate that the G899D Upc2A is no longer sensitive to inhibition by ergosterol binding and instead localizes to the nucleus. The elevated nuclear levels of G899D Upc2A are sufficient to relieve any requirement for the increased activity that would normally be provided by Ypk2 and suppressed the increased FLC susceptibility normally seen in *ypk2Δ* strains.

These data also suggest that Ypk2 may act to coordinate the biosynthesis of sphingolipids and sterols, the two main components of lipid rafts (reviewed in (SIMONS AND SAMPAIO 2011)). In *S. cerevisiae*, Ypk kinases stimulate the activity of the serine palmitoytransferase enzyme, the first committed step in sphingolipid biosynthesis (BRESLOW *et al*. 2010; ROELANTS *et al*. 2011). The ScYpk kinases have been found to regulate the retrograde movement of ergosterol from the plasma membrane to the ER (MURLEY *et al*. 2015; MURLEY *et al*. 2017; ROELANTS *et al*. 2018) but have not been reported to influence the biosynthesis of this important membrane lipid. Our finding that Ypk2 activity stimulates expression of several different ERG genes indicates that this kinase stimulates production of enzymes that ultimately produce ergosterol. While much less is known of the role of Ypk2 (or Ypk1) in regulation of sphingolipid biosynthesis in *C. glabrata*, the myriocin hypersusceptible phenotype shown here is consistent with Ypk2 acting as it does in *S. cerevisiae* to stimulate sphingolipid biosynthesis. Together, these data implicate Ypk2 as serving to link production of the two main components of lipid rafts: ergosterol and sphingolipids.

The role of Ypk2 as a determinant in susceptibility to three different antifungal drugs is potentially due to the effect of this kinase on membrane biology, illustrated by its impact on ergosterol and sphingolipid biosynthesis. Altering levels of important membrane lipids, that are estimated to make up roughly 50% of the plasma membrane when considered together (SCHNEITER *et al*. 1999; KODEDOVA *et al*. 2019), have been linked with changes in membrane trafficking of proteins as well as activity levels of proteins embedded in these membranes. For example, changes in membrane ergosterol levels are well-established to impact delivery of transporter proteins to the plasma membrane as well as to influence targeting of ER resident proteins. Membrane proteins that may act to efflux drugs from the cell (such as Cdr1) are likely to be impacted by changes in the membrane environment in which it functions. Additionally, the presumptive echinocandin target enzyme, β-1,3-glucan synthase, is also a plasma membrane resident protein. Its trafficking and activity will also be a function of its membrane milieu. We suggest that changes in the lipid composition of the plasma membrane, responsive to Ypk2, may be crucial in the determination of susceptibility to multiple antifungal drugs.

Finally, signaling pathways are well-established to interact. This was recently illustrated in *S. cerevisiae* by experiments demonstrating that activation of the cell wall integrity pathway negatively impacts TORC2 signaling and can block Ypk kinase activation (NOMURA AND INOUE 2024). This signaling interaction can lead to increased susceptibility to sphingolipid biosynthesis inhibitors and hints at the potential complexity underlying the phenotypic effects we have seen elicited by loss of Ypk2 in *C. glabrata*. Experiments directly examining the behavior of specific members of signaling pathways are required to reveal the molecular explanations for observed phenotypes described here.

## Acknowledgements

This work was supported by NIH AI152494 and the German Research Foundation (DFG) through the TRR 124 FungiNet, project number 210879364, projects A1 and Z2. We thank Drs. Damian Krysan and Scott Filler for valuable discussions during the course of this work. We are grateful to Olaf Kniemeyer for his critical reading of the manuscript.

## Supplementary Figure

**Supplementary Figure 1.**
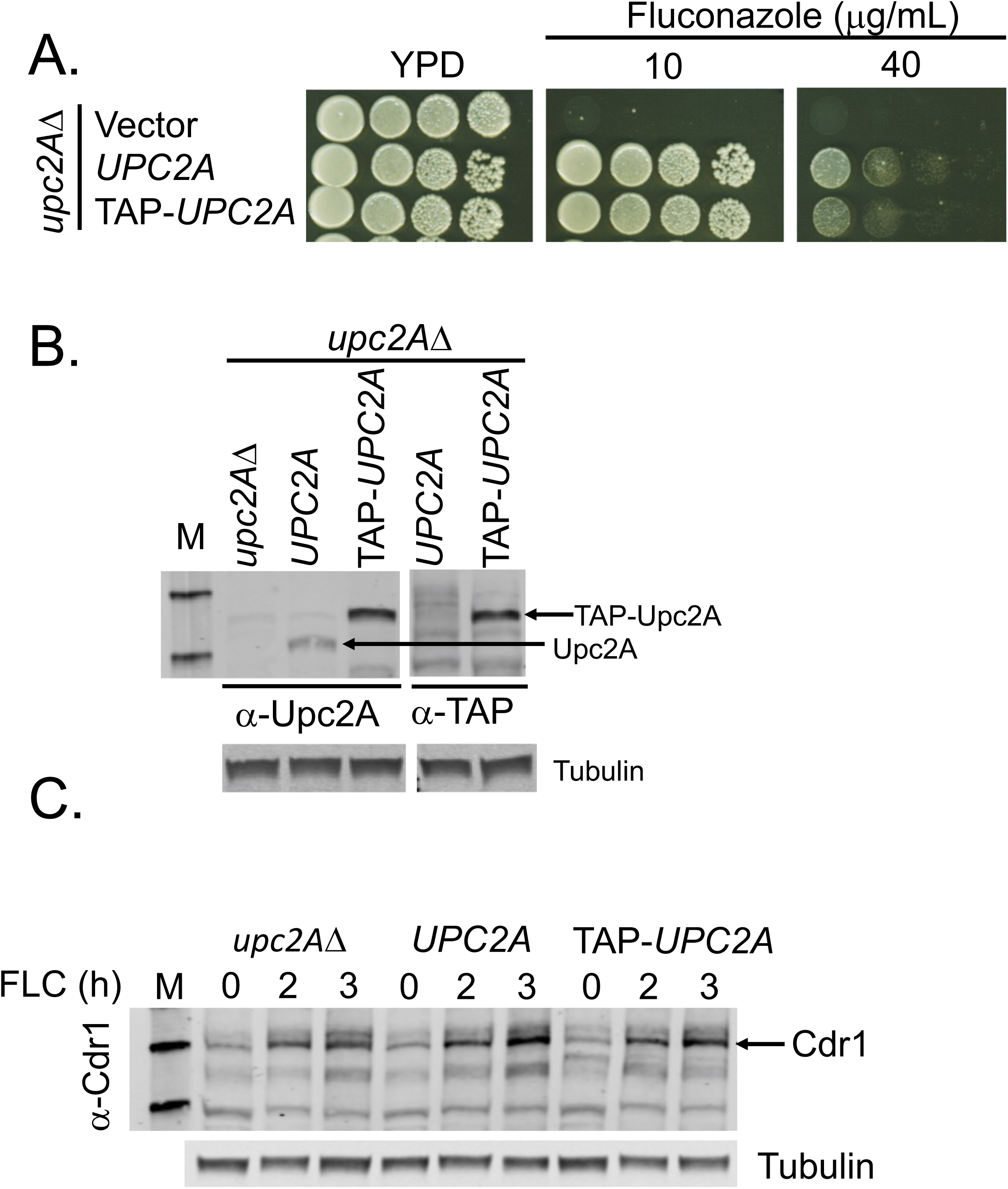
Characterization of the TAP-*UPC2A* strain. Fluconazole susceptibility of the TAP-UPC2A is similar to untagged strain. Western blot detection of TAP-Upc2A protein levels by α-Upc2A or α-TAP antibodies. Western blot detection of induction of Cdr1 by α-Cdr1 antibody in the *upc2A*Δ, wild type *UPC2A* and TAP-tagged *UPC2A* expressing cells untreated (-) or treated with fluconazole (20 ug/mL) for 2 or 3 hours. TAP purification of Upc2A from sample untreated (-) or treated (+) with 20 ug/mL of fluconazole for 2 hours for the mass spectrometry analysis. Input corresponds to the Upc2A present in the 1% fraction of the whole cell lysate. Final elution indicates a concentration of the purified Upc2A after two steps of TAP-purification. An aliquot of the same concentration of the purified Upc2A as detected by the western blot in the final elution was sent for the mass-spectrometry.

